# Natural variants of von Willebrand factor R1205 causing von Willebrand disease with accelerated von Willebrand factor clearance: *in silico* docking models and energetics of the interaction with both LRP1 and GpIb A1 domain

**DOI:** 10.1101/2025.08.22.671727

**Authors:** Monica Sacco, Stefano Lancellotti, Antonietta Ferretti, Maria Basso, Leonardo Di Gennaro, Giancarlo Castaman, Raimondo De Cristofaro

## Abstract

Type 1 von Willebrand disease (VWD) is often caused by variants in von Willebrand factor (VWF), including p.R1205H (“Vicenza mutation”), which accelerate VWF clearance via macrophage receptor LRP1 and impair platelet adhesion. However, the structural mechanisms underlying these phenotypes remain unclear. Here, we use integrative computational modeling (I-TASSER, HADDOCK2.4, and PRODIGY) to predict how p.R1205H/C/L/S variants alter VWF interaction with LRP1 and GPIbα. Our models reveal that R1205 acts as a structural hinge: its variants disrupt polar networks in VWF’s D3 domain, exposing neo-epitopes that enhance LRP1 binding (ΔΔG up to −5.3 kcal/mol) while destabilizing the A1 domain’s α1-β2 loop, reducing GPIbα affinity (33-fold for R1205L/C). These findings explain clinical observations, p.R1205H rapid clearance yet retained platelet adhesion, and establish R1205 as a dual-functional switch regulating VWF circulatory lifetime and hemostatic activity. This analytical procedute provides a template for predicting pathogenicity of VWF variants and designing targeted therapies for VWD.

**Author Summary:** In our study, we explored how specific genetic changes in von Willebrand factor (VWF), a protein crucial for haemostsis, lead to a bleeding disorder called von Willebrand disease (VWD). We focused on mutations at position R1205 in VWF, known to cause rapid clearance of the protein from the bloodstream, resulting in low levels and impaired clotting. Using advanced computer modeling, we investigated how these mutations alter VWF’s interactions with two key partners: LRP1, a receptor that removes VWF from circulation, and GpIbα, a platelet receptor essential for clotting. Our models revealed that R1205 acts like a structural hinge. Mutations disrupt its stabilizing role, exposing regions that bind more tightly to LRP1, thus explaining the accelerated clearance seen in patients. Simultaneously, these changes can also weaken VWF’s ability to interact with platelets, further impairing platelet haemostatic function. This work provides a blueprint for understanding how some VWF variants cause disease and could guide the design of targeted therapies to correct or compensate for these defects, improving treatment for VWD patients.

## Introduction

The high molecular weight glycoprotein VWF is essential to maintain normal hemostasis through its ability to mediate platelet-subendothelial interactions as well as interaction with platelets to support their aggregation at the site of damaged endothelium. The type 1 von Willebrand disease (VWD1) shows a variable phenotype associated with a mild to moderate quantitative reduction of VWF levels, whose concentrations range from 5% to 50% of normal [1]. This disease derives from several types of alterations including defective RNA or protein synthesis, early protein degradation inside the cell, ineffective secretion, accelerated plasma clearance, or mutations that result in a null allele [2]. The disease referred to as “Vicenza VWD”, is a typical subtype of type 1 VWD due to the p.R1205H missense variant, whereby a relatively severe phenotype is associated with an accelerated VWF clearance [3, 4]. Hence, the Vicenza variant causes a significant reduction in VWF antigen (VWF:Ag) and VWF Ristocetin Cofactor Activity (VWF:RCo) ≤0.20 U/mL, as well as Factor VIII levels <0.30 U/mL At variance, platelet VWF levels and function are normal [3, 5–7]. The variable genetic and protein phenotypes is reflected by a wide range of bleeding scores, ranging from 2 to 17, (mean value=8) [1, 8, 9]. Other types 1 VWD variants occurring at the same place, such as p.R1205C, p.R1205L, and p.R1205S share similar characteristics with VWD Vicenza, being also characterized by accelerated clearance. This pathogenetic mechanism was demonstrated via desmopressin (DDAVP) studies [2] and human recombinant protein infusion in the VWF knockout mouse [10], as well as through high VWFpp/VWF:Ag ratios, observed in patients’ plasma [5, 11]. The p.R1205H missense variant occurs in the E3 module of the VWF D3 domain [2]. This pathological variant in some cases shows even alterations in the typical multimer triplet band pattern in SDS-agarose gels with a predominance of ultra-large VWF multimers [5, 12], Also these changes were attributed to the very rapid clearance of VWF multimers [4, 11, 13] that limit their physiological proteolytic processing by ADAMTS-13 [14, 15]. Following DDAVP infusion, plasma VWF:Ag levels increase significantly in VWD-Vicenza individuals [12, 14]. However, the half-life of the secreted VWF-R1205H is markedly reduced compared with the wild-type (WT) VWF [2, 4, 6]. The direct role of the arginine residue at position 1205 of the mature VWF molecule in regulating the half-life of the protein was confirmed in several studies using VWF -/- mice. The mean half-life of human p.R1205H VWF was markedly shorter than that of human WT-VWF in VWF -/- mice [10]. Likewise, the half-life of murine p.R1205H VWF was also significantly short following intravenous injection in VWF deficient mice [16]. As anticipated above, other missense variants of R1205 in the D3 domain of VWF are associated with low VWF levels and possibly accelerated clearance from circulation: 1) p.R1205C; 2) p.R1205L; and 3) p.R1205S [3, 17], although a direct demonstration is lacking. Furthermore, a conformational linkage between the A1 and D3 domains of von Willebrand factor (VWF) has been studied in the context of VWF’s structure and function [18–24]. The A1 domain is responsible for binding to platelet glycoprotein Ib (GPIb), while the D3 domain is involved in the regulation of VWF multimerization, stability, and FVIII interaction. Specifically, the D3 domain plays a regulatory role by influencing the conformation of the A1 domain, possibly modulating its ability to bind GPIbα [25–27].

These findings strongly suggest that R1205 would play a relevant structural and functional role, modulating the interaction of VWF with the various scavenging receptors and possibly with the platelet receptor GPIbα. Hence, this study was aimed at investigating by molecular modeling the structural and functional consequences of the p.R1205H, p.R1205S, p.R1205C, and p.R1205L VWF variants on the interaction with both domain IV of the macrophage LRP1 and the N-terminal domain of the platelet GpIbα.

## Results

### Molecular modeling of WT and R1205 pathological variants of VWF

Table 1 lists the TM and RMSD values for both the WT and VWF variants. All 3D models had a template modeling score (TM-score) >0.50, indicating acceptable structural topology. Only the p.R1205C variant exhibited a TM-score at the limit of structural acceptability. Overall, the topology of the VWF forms is roughly characterized by three globular parts (764-1180, 1211-1866, and 1920-2191). These globular parts are connected by two linker regions (1181-1210 and 1867-1919), as shown in Figure 1. As to the R1205X variants, analysis of the molecular modeling results showed that the substitution of the Arg with different amino acids alters the conformation of discrete regions of the molecule, as, for instance, in the case of p.R1205H (see Fig. 2). However, the TM-align program provided in all cases a TM-score value ranging from 0.64 (p.R1205L) and 0.84 (p.R1205C), with intermediate values for p.R1205H (0.76) and p.R1205S (0.75). These values show that all these VWF mutants have roughly the same overall structure [28].

**Figure 1.**
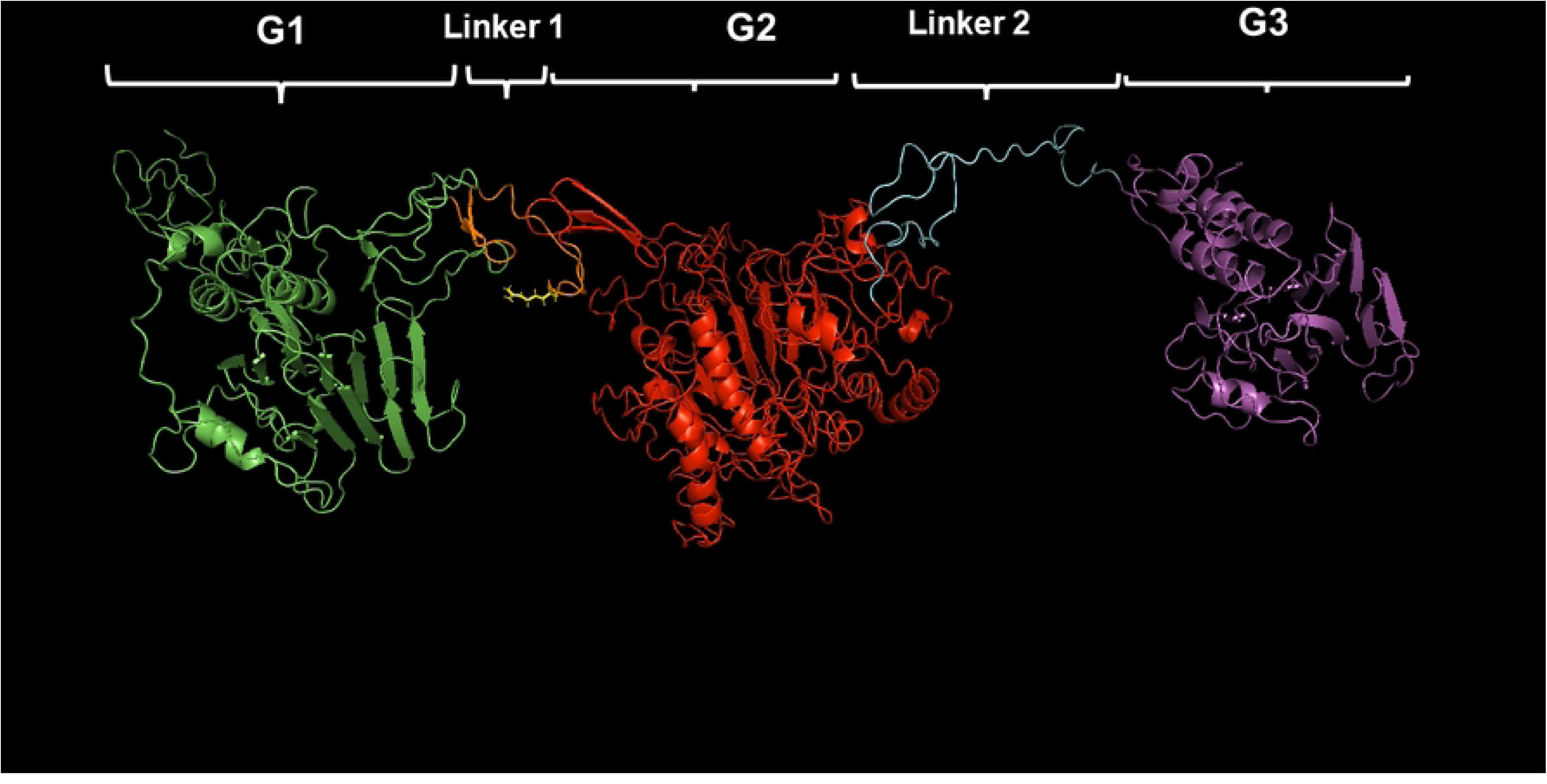
Best model of the 764-2191 region of WT-VWF. The three globular regions (G1-G2-G3) are shown in green, red, and magenta, respectively, while the linker region 1 (1181-1210) is shown in orange, and the second linker region (1867-1919) is shown in deep blue. The side chain of R1205 is shown in yellow sticks. The model was obtained with the I-TASSER program, whereas the manipulation was accomplished with Pymol.

**Figure 2.**
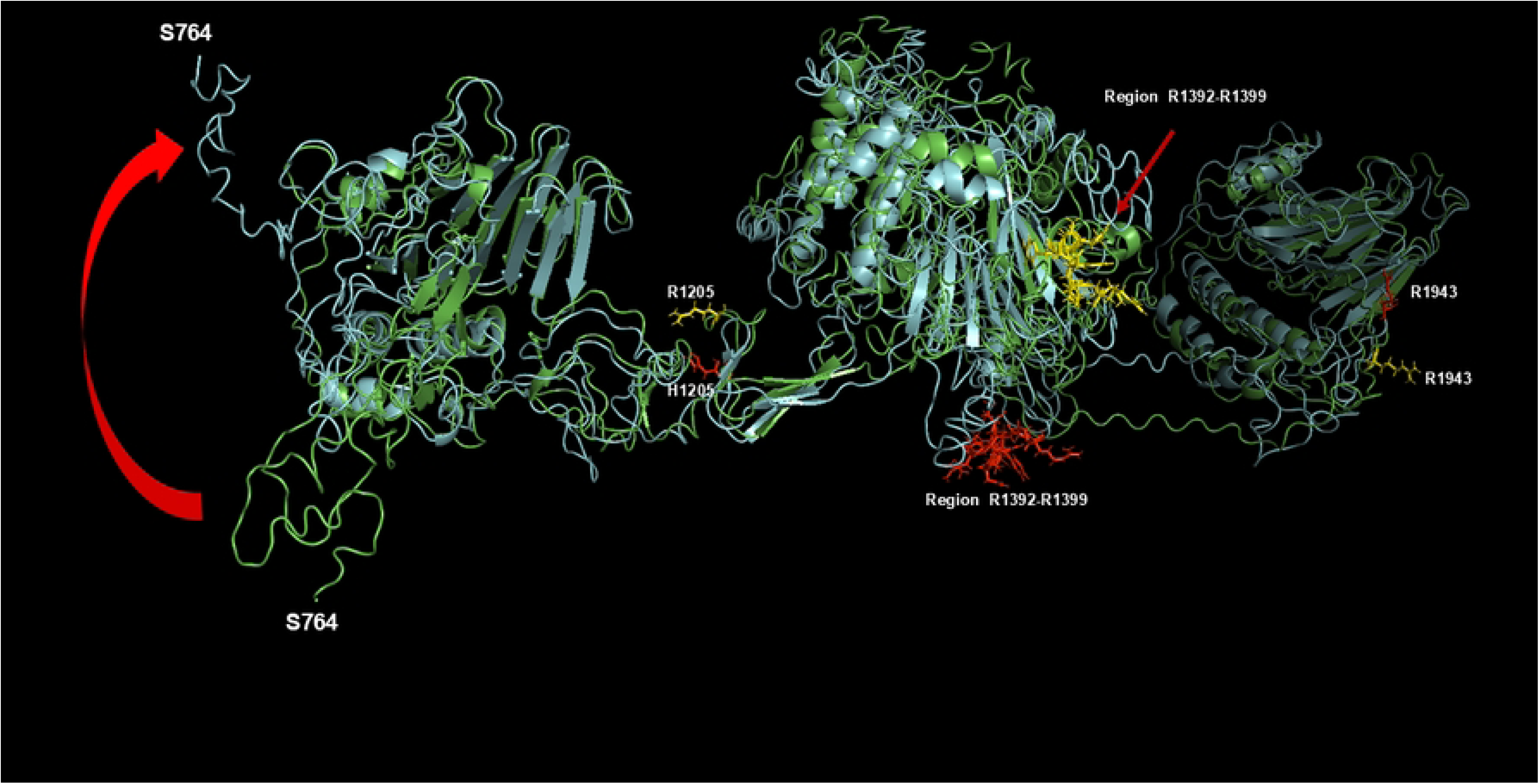
Superposition of WT-VWF (green ribbon) and p.R1205H (cyan ribbon). The region R1392-R1399 and R1943, involved in binding to LRP1 domain IV, are shown as yellow sticks and red sticks for WT and p.R1205H VWF, respectively. The R1943 side chains of both WT (yellow sticks) and p.R1205H variant (red sticks) are also shown. Note the large topological change for the region S764-T789, likely deriving from a huge conformational flexibility (curve red arrow). The superposition and alignment of the two structures were performed using the PyMOL program.

**Table 1.** Results of the I-Tasser analysis concerning the 3D models of the monomer WT (764-2191) and R1205 VWF variants.

| Species | Estimated TM-score | Estimated RMSD (Å) |
| --- | --- | --- |
| WT | 0.53±0.14 | 10.1±4.6 |
| p.R1205H | 0.52±0.11 | 10.5±4.8 |
| p.R1205S | 0.56±0.15 | 15.4±3.4 |
| p.R1205C | 0.45±0.15 | 15.5±3.3 |
| p.R1205L | 0.81±0.09 | 8.0±4.4 |

In the WT-VWF molecule, R1205 forms a series of polar and salt bridges at a distance ≤3 Å with A1207, E1185 and E1158 (see Fig. 3). This setting contributes to stabilizing a correct topology of the region between the G1 and Linker 1 parts. The lack or change of these bonds in the other R1205X variants allosterically induces the rearrangement of regions involving both the segment R1392-R1399 and R1943, involved in binding to LRP1 domain IV (see Figure 4A). The relevance of the positive charge at VWF-R1205 has been recently remarked by the results obtained with SPR experiments on WT VWF-LRP1 interaction by Atiq et al. [4]. Hence, the conformational effects arising from substitution of R1205 with different amino acids would determine the functional changes in the interaction with LRP1 (see below). The changed polar interactions of the side chains of H1205, C1205, L1205, and S1205 with surrounding residues are shown in Supplementary Figures S1-S4. In these cases, the lack of interactions of the side chain of R1205 with surrounding amino acids causes long-range conformational changes, as shown in Figure S5.

**Figure 3.**
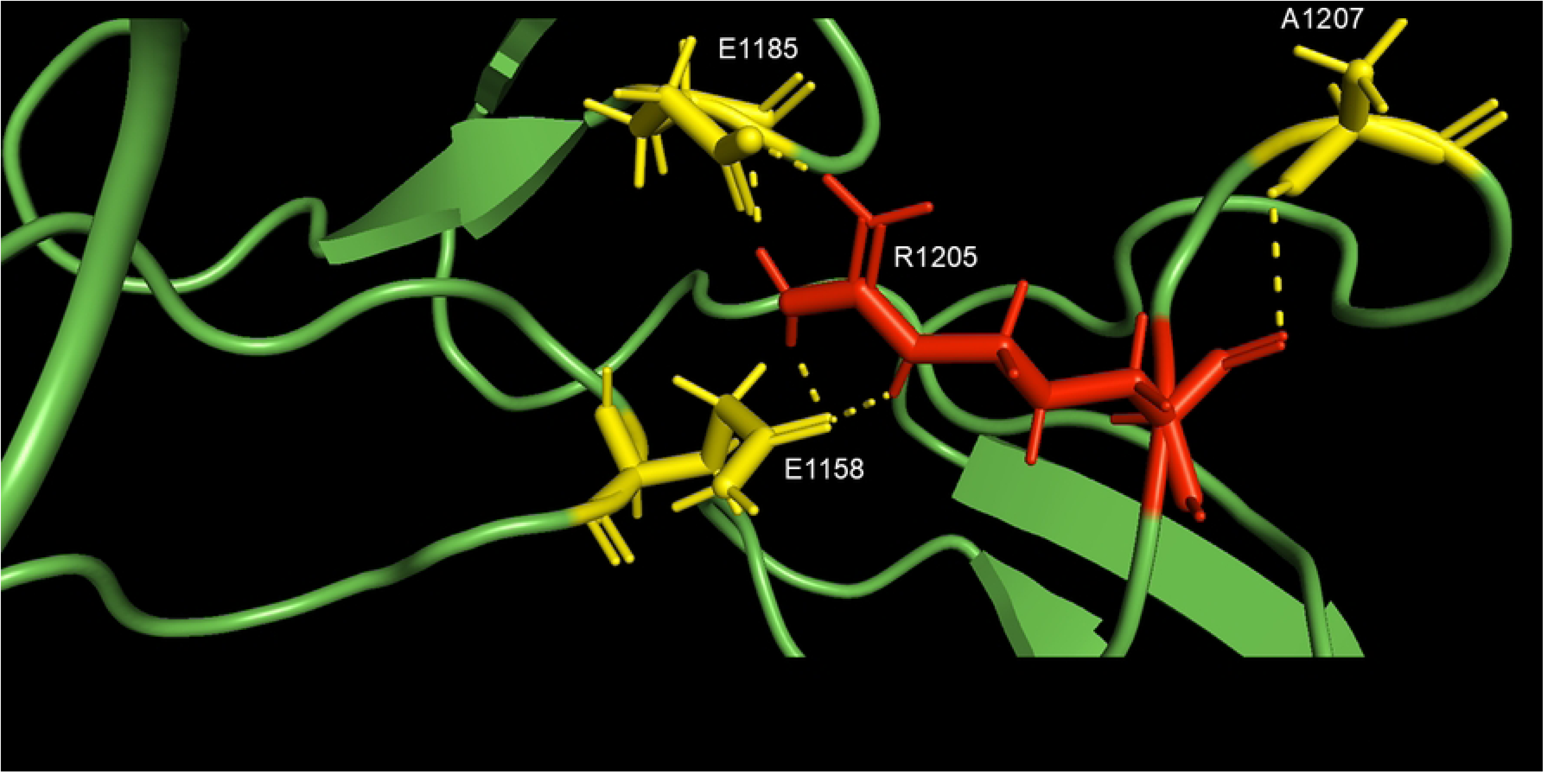
Polar interactions ≤3 Å (dashed lines) between the side chain of R1205 (shown in red sticks) in the WT form of VWF and the surrounding amino acids (yellow sticks). The model was obtained with the I-TASSER program, whereas the manipulation was accomplished with Pymol.

**Figure 4.**
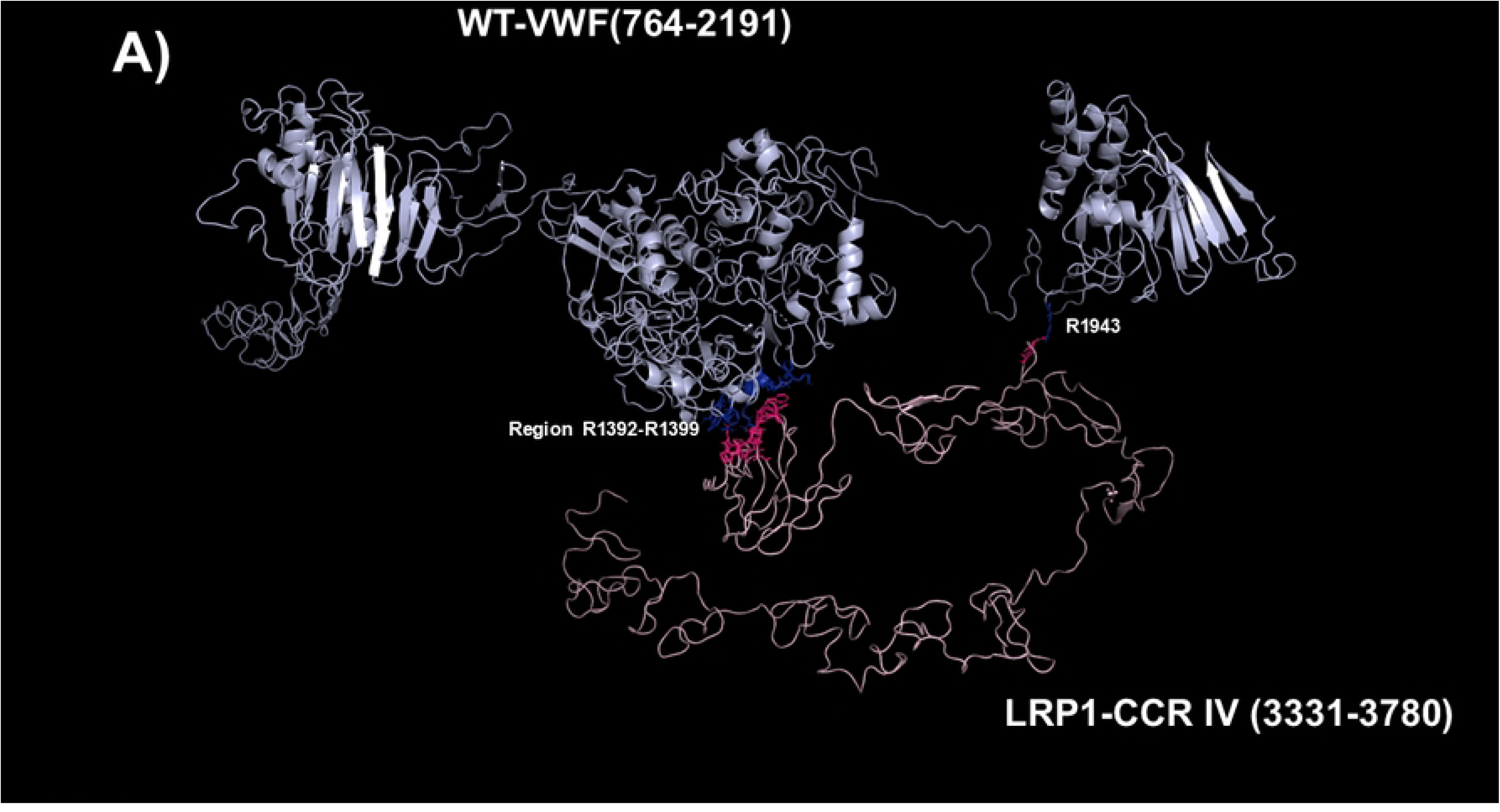
Molecular models obtained with I-TASSER and Haddock 2.4 programs of the adducts between with A) WT-VWF; B) p.R1205H-VWF; C) p.R1205C-VWF; D) p.R1205LVWF; E) p.R1205S-VWF. The VWF constructs are shown in cyan, whereas the CCR IV of LRP1 is shown in pink. The interacting residues of each complex are listed in Table 2. The interacting side chains of VWF constructs are shown as blue sticks, while those pertaining to LRP1 are shown as warmpink sticks.

**Table 2.** Interface features of the adducts between WT and VWF variants at R1205 and LRP1 domain IV. Chain A represents VWF, whereas Chain B is LRP1 domain IV.

|  |  |  |  |  |  |  |  |
| --- | --- | --- | --- | --- | --- | --- | --- |
| **VWF species** | **Chain** | **No. of Interface residues** | **Interface Area (Å<sup>2</sup>)** | **No. of Salt bridges** | **No. of disulphide bonds** | **No. of hydrogen bonds** | **No. of Non-bonded contacts** |
| **WT** | **A** | 9 | 565 | 7 |  | 12 | 78 |
|  | **B** | 11 | 590 |  |  |  |  |
| **p.R1205H** | **A** | 23 | 901 | 6 |  | 14 | 127 |
|  | **B** | 20 | 1015 |  |  |  |  |
| **p.R1205C** | **A** | 21 | 1268 | 2 |  | 6 | 82 |
|  | **B** | 23 | 1265 |  |  |  |  |
| **p.R1205L** | **A** | 14 | 990 | 6 |  | 12 | 102 |
|  | **B** | 21 | 907 |  |  |  |  |
| **p.R1205S** | **A** | 21 | 1354 | 1 |  | 8 | 77 |
|  | **B** | 23 | 1345 |  |  |  |  |

The same computational methodology described above was used to predict the ab initio structure of LRP1 receptor cluster IV (V331–P3780).The best final model is shown in the Supplementary Figure S6.

All PDB files corresponding to the obtained structural models are available in the Supporting Information file..

### Models of the complexes between LRP1 Cluster IV and the VWF species

As paradigmatic adducts between LRP1 cluster IV and the VWF models, we selected those characterized by higher negative HADDOCK and Z-scores. Figure 4A-E shows the best models obtained using the HADDOCK platform. In all the adducts but p.R1205L-VWF and p.R1205S, G2 and G3 domain of VWF are engaged in the interaction. Instead, in p.R1205L and p.R1205S adducts, G1 and G2 domains participate in the interaction with LRP1 and change the energetics of binding. Table S1 lists all the hydrogen bonds and salt bridges present in all the complexes formed by all the VWF forms and LRP1. WT-VWF interacts with the sequence 1392-1399, along with R1943. All VWF mutants except p.R1205C also engage R1399 for binding to LRP1, though they utilize additional regions due to conformational changes induced by the mutations (see Table S1 for details). Table 2 shows that the VWF interaction area for LRP1 binding is significantly larger for the R1205 variants than for the WT form, changing from about 570 Å2 up to 1350 Å2 in p.R1205S mutant. Because of the conformational effects, there is a strong increase of the predicted interaction affinity of the VWF variants compared to the WT form. The binding affinity of different VWF forms exhibits a linear dependence on the interface area, as shown in Figure 5. The ΔG of equilibrium binding constants of both WT- and p.R1205X variants are reported in Table 3 and visually shown in Figure S7.

**Figure 5.**
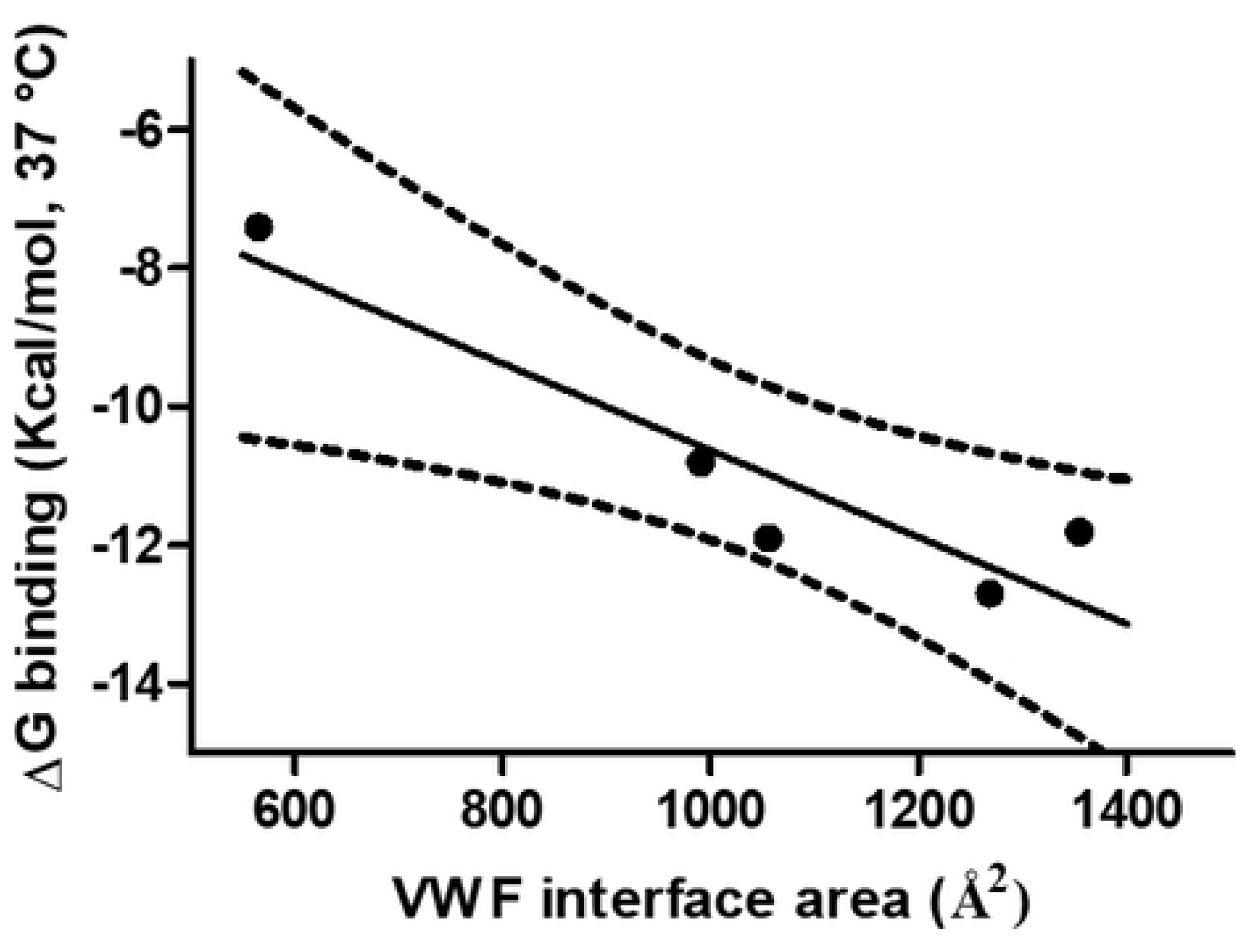
Values of ΔG of binding of the VWF forms to LRP1 as a function of VWF interface areas. The slope of the linear regression is −0.006271 ± 0.001458 (p= 0.0231).

**Table 3.**
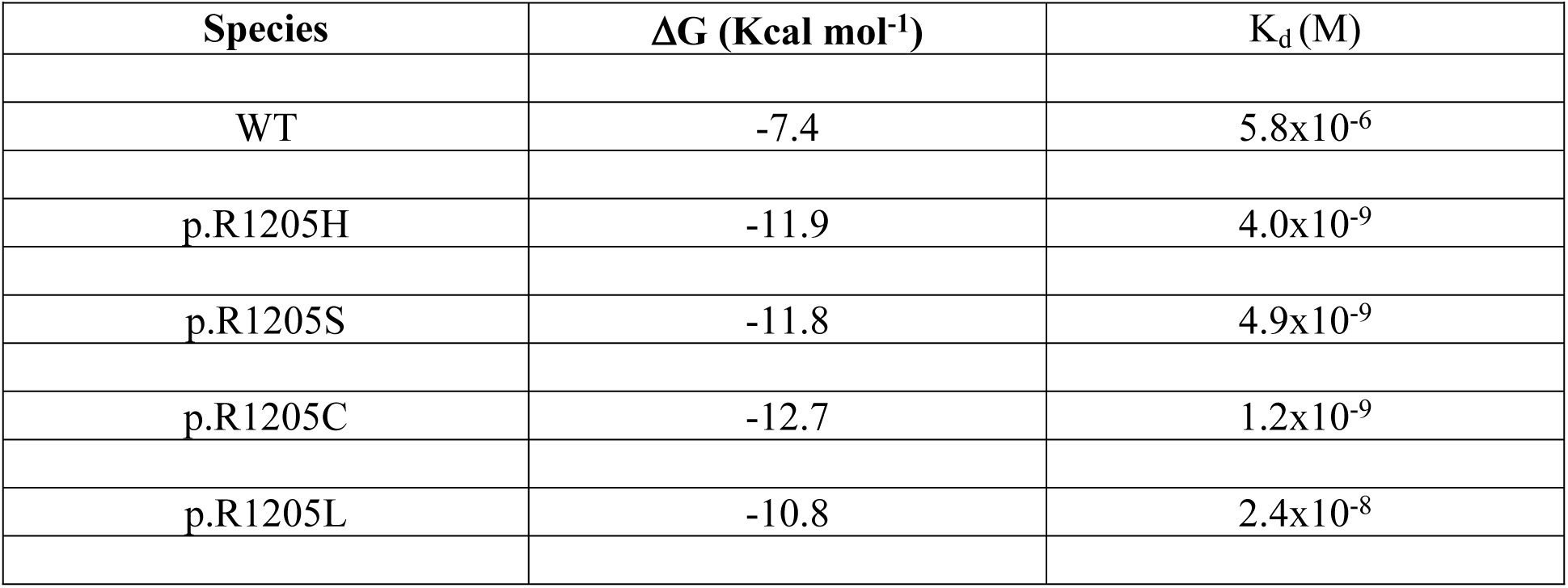
Calculated ΔG of binding of different WT monomer species and LRP1-Cluster IV with the corresponding K_d_ values calculated by the PRODIGY program (T=37 °C)

The PDB files for all modeled structures of WT and variant VW with LRP1 are included in the Supporting Information

### Models of the complexes between N-terminal domain of platelet GpIbα and the various VWF species

The deposited pdb file 1SQ0 was considered as a template for the analysis of the models of the complexes between the N-terminal portion of GpIbα (H1-T266) and the VWF(764-2191) forms. In the pdb file 1SQ0 concerning the interaction between N-Terminal GpIbα and the A1 domain of VWF several hydrogen bonds are formed between S1325, A1327, R1334, E1359, K1362, F1366, Q1367, and R1395 of VWF and M239, K237, D18, S39, Y238, P198, N226, Y228, K152, T175, and E225 of GpIbα, respectively. Thus, an extensive region of the G2 domain of the VWF(764-2191) molecule is involved in this interaction. Overall, this setting is also followed by the R1205X variants (see Figure 6A-B). All p.R1205X variants exhibit reduced binding affinity for GpIbα. Compared to WT-VWF, p.R1205H displayed approximately one-third the binding affinity, while p.R1205L and p.R1205C showed even weaker binding. The p.R1205S variant displayed intermediate affinity (Table 4). A detailed inspection of the models of both p.R1205L and p.R1205C variants shows that the conformation of the α1-β2 loop of the VWF A1 domain (E1294-D1323, G2 in the model shown in Figure 1), which plays a major role in modulating the affinity of VWF for GpIb, is largely altered in comparison to the WT form, dampening the affinity for the platelet receptor, as shown in Figures 7-8.

**Figure 6.**
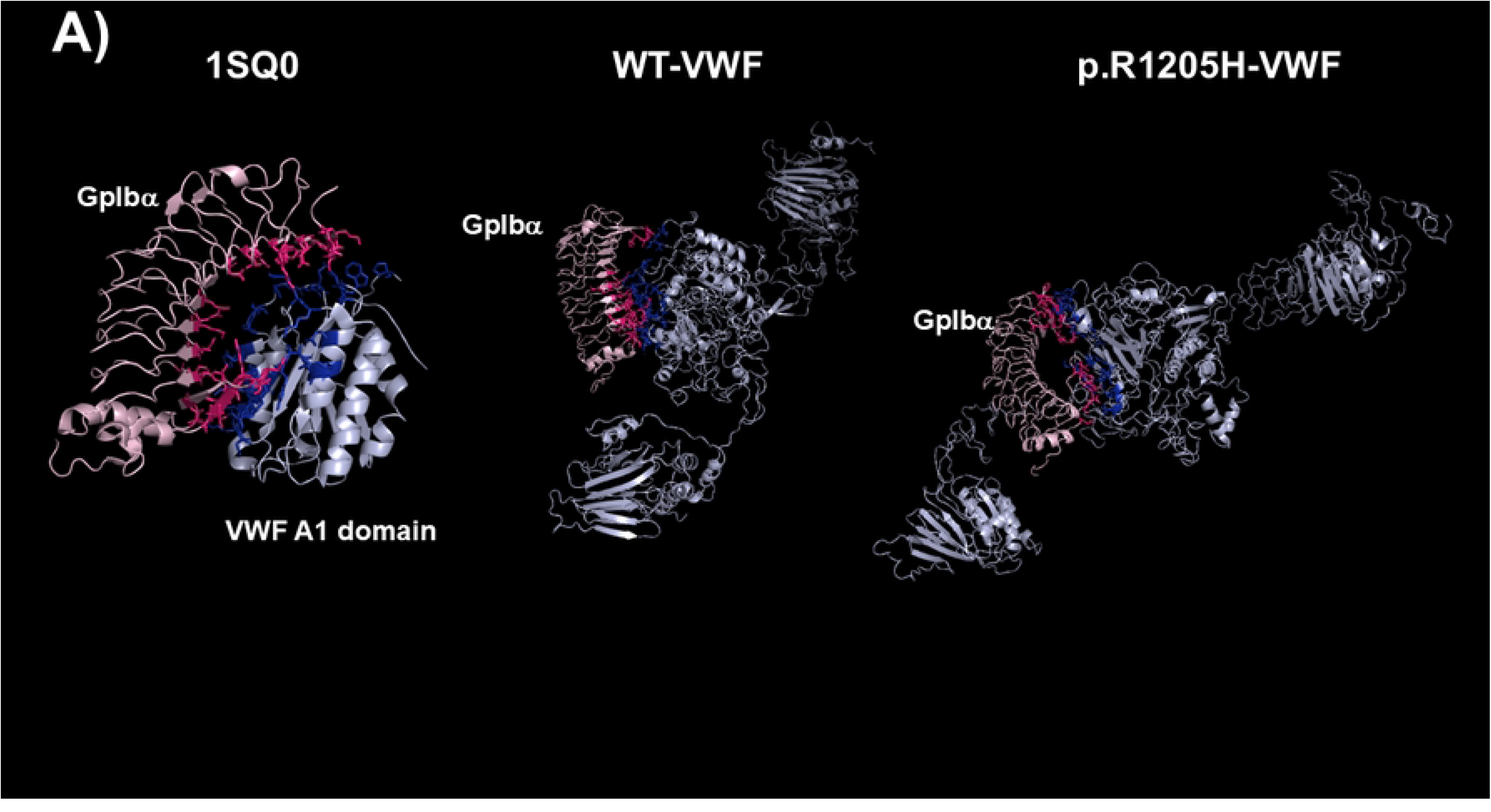
**A)** Best models of the 764-2191 region of WT-VWF and R1205H-VWF models bound to the N-terminal (1-266) domain of platelet GpIbα. For comparison, the crystal structure of the N-terminal (1-266) region of GpIbα bound to the VWF A1 domain (pdb: 1SQ0) is also shown. **B)** The models of the complexes of the other p.R1205X variants (p.R1205C, p.R1205L, and p.R1205S) with the same GpIb domain. In all cases, GpIbα binds to the G2 region of VWF. The models of VWF(764-2191) are shown cyan, while the GpIbα molecule is shown in pink. The side chain of interacting residues of GpIbα is shown in warm pink, whereas those of VWF are shown as blue sticks. The models were generated by the HADDOCK program, whereas the rendering was accomplished with Pymol.

**Figure 7.**
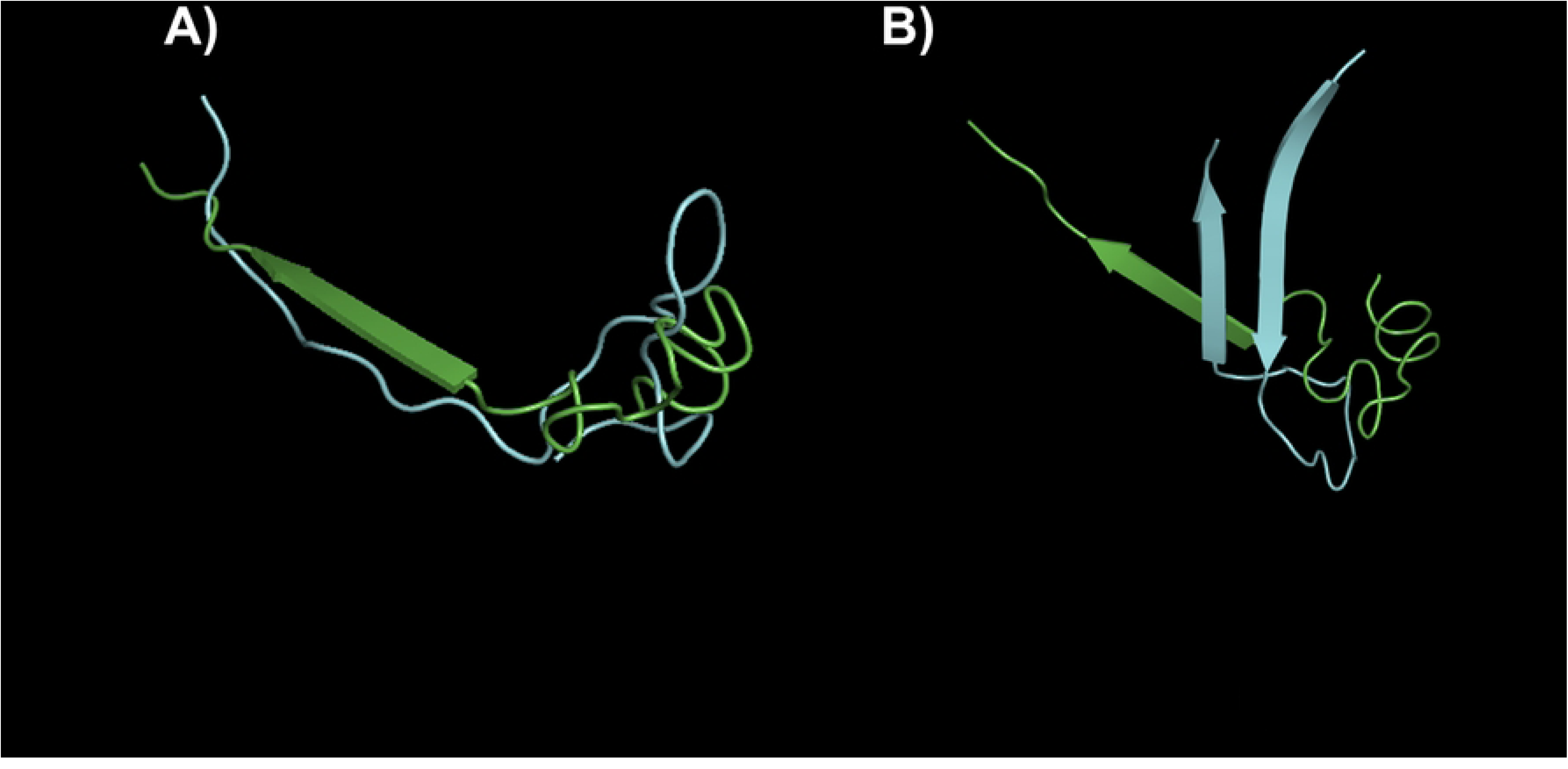
**A)** The best model of the α1-β2 loop (E1294-D1323) of the WT-VWF(764-2191) shown in green superposed to the same region of p.R1205L and **B)** p.R1205C both shown in cyan. The models were generated by the I-TASSER program, whereas the rendering was accomplished with Pymol.

**Figure 8.**
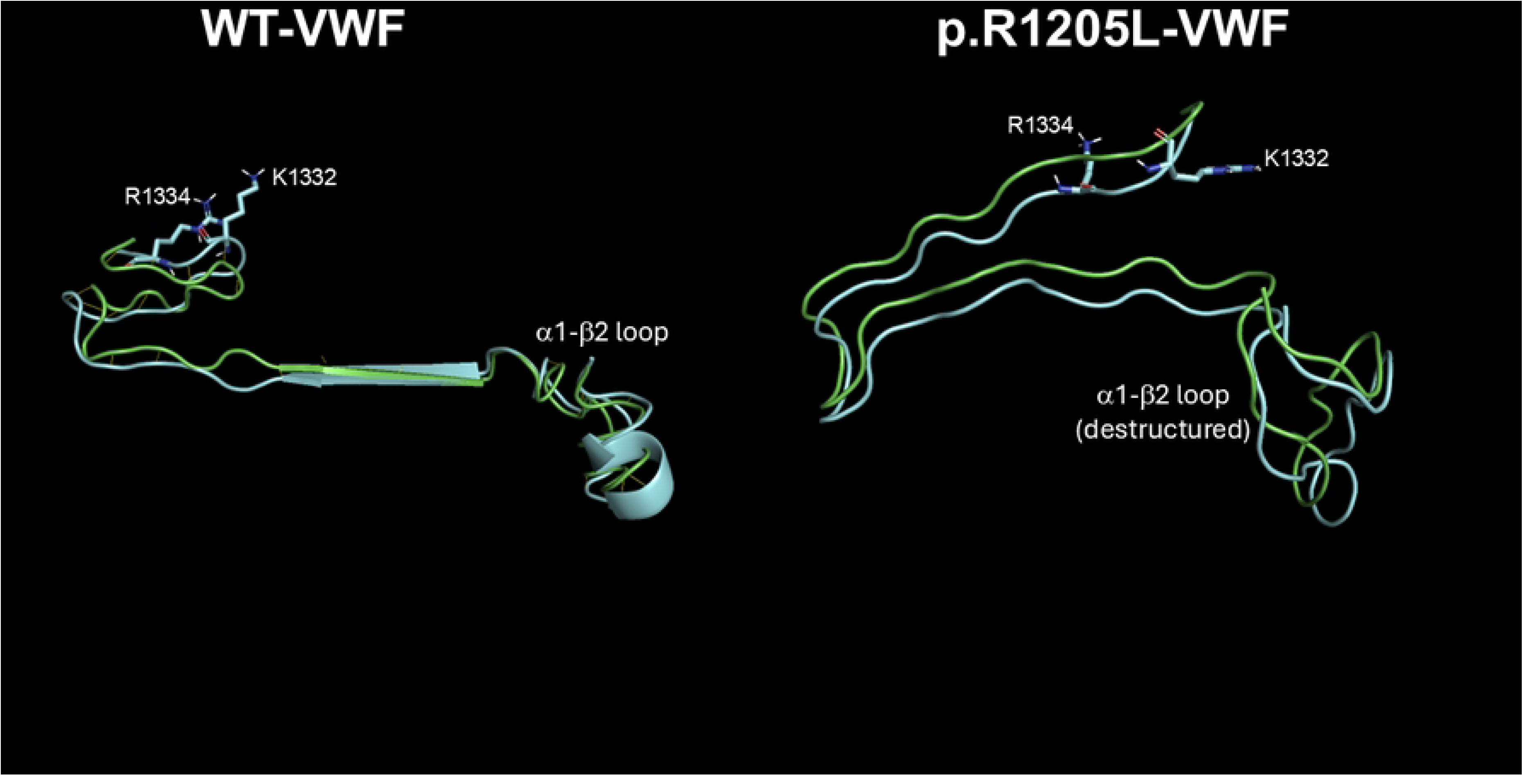
Movement of region 1294–1336 in both WT-VWF and p.R1205L VWF from the GpIb-free conformation (green) to the GpIb-bound conformation (cyan). This region includes the α1-β2 loop (1309–1314), which undergoes a significant conformational change, flipping by ∼1.5 Å upon binding to GpIbα. This allows other residues, such as R1334 and K1332, to directly contact K237 and E225 of GpIbα (not shown). Note the altered orientation of K1332 and R1334 in the VWF variant, which makes interaction with GpIbα more difficult. Similar findings were observed with the p.R1205C-VWF variant. Images were rendered using PyMOL.

**Table 4.** Calculated ΔG of binding and the corresponding K_d_ values for the different VWF species and the N-terminal (H1-T266) domain of platelet GpIbα. The binding parameters were calculated by the PRODIGY program at T=37 °C.

| Species | $\Delta G$ (Kcal mol<sup>-1</sup>) | $K_d$ (M) |
| --- | --- | --- |
| WT | -11.7 | $5.6 \times 10^{-9}$ |
| p.R1205H | -11.1 | $1.5 \times 10^{-8}$ |
| p.R1205S | -10.2 | $6.0 \times 10^{-8}$ |
| p.R1205C | -9.5 | $1.9 \times 10^{-7}$ |
| p.R1205L | -9.4 | $2.2 \times 10^{-7}$ |
| 1SQ0* | -10.1 | $7.2 \times 10^{-8}$ |
| * PDB file of the complex formed by A1 VWF domain and N-Terminal part of GpIbα | | |

The PDB files for all obtained VWF-GpIb structural models are included in the Supporting Information.

## Discussion

The study addresses a clinically significant question, namely how variants VWF at R1205 affect its interaction with both LRP1 and GPIbα, leading to VWD1, functionally characterized by normal or slightly prolonged bleeding time in vivo and strongly altered Platelet Function Analyzer (PFA) test in vitro [2, 5, 29–38]. Macrophage LRP1 mostly modulates the clearance of VWF and its complex with FVIII. LRP1 would accomplish this function via direct shear stress-induced interactions with VWF while blood containing VWF is flowing along LRP1-expressing macrophages, or as part of a functional LRP1–β_2_-integrin complex, as previously demonstrated [39]. However, although this represents the predominant clearing pathway of VWF, it is not the only mechanism involved in the VWF clearance from circulation. Other systems, in fact, contribute to this function such as the asialoglycoprotein receptor, CLEC4M, or members of the Siglec family [13].

The present findings agree with the clinical phenotype of the “Vicenza” type 1C VWD, characterized by very rapid clearance of the p.R1205H variant. Previous studies demonstrated that in VWF-null mice, recombinant VWF with the p.R1205H mutation has a ≈7-fold shorter plasma residence time (0.3 h) than wild-type VWF (2.8 h). Of interest, in that study the clearance was independent of multimeric size [17]. This aligns with our finding that the p.R1205H monomer model shows much higher LRP1 affinity than WT, as also recently found in binding experiments with surface plasmon resonance [4]. Liver and spleen macrophages likely regulate VWF clearance by binding and endocytosing VWF multimers, as supported by experimental data [17, 39]. Macrophage depletion also significantly prolonged the survival of p.R1205H, p.R1205S and p.R1205C VWF species [17]. Notably, macrophage depletion inhibited p.R1205H VWF clearance to a greater degree than that observed with WT-VWF, suggesting that macrophage-mediated clearance is the predominant pathway through which p.R1205H is cleared from circulation. Hence, this very high affinity for macrophage LRP1 would represent the main mechanism playing a prominent role in clearance of the VWF–FVIII complex [17]. The molecular details of the VWF regions involved in the clearance mechanism have not yet been fully defined. However, the knowledge of the functional behavior of the natural mutants associated with enhanced VWF clearance unequivocally demonstrated that the involved amino acid residues are clustered within the D’D3 and A1A2A3 domains [40]. Experimental studies demonstrated in fact that recombinant A1A2A3 domains infused in mice were cleared from circulation at a similar rate to full-length VWF [13]. Notably, the fragment D’D3A1A2A3 was cleared more slowly, suggesting that the D’D3 domains may have a regulatory role in VWF clearance, slowing the process of interaction with VWF scavenger receptors [13]. A similar role of the D’D3 domain was demonstrated for the A1-VWF domain interaction with the platelet receptor GpIb-IX-V [26]. Any variations that can perturb the native conformation of the D3 domain around R1205 can abrogate the inhibitory effect of the D’D3 domain. The results obtained in the present molecular modeling investigation concerning p.R1205H, p.R1205S, p.R1205L, and p.R1205C fully agree with this scenario. While pathological variants retain WT-VWF overall fold, local structural perturbations, caused by R1205 variants, alter binding to the LRP1 domain IV. A critical question is how these variants enhance VWF affinity for macrophage LRP1. In WT-VWF, R1205 establishes both polar contacts (≤3 Å) and salt bridges with A1207, E1185, and E1158 (see Fig. 2). Functioning as a structural hinge, these interactions maintain proper spacing between VWF’s G1, G2, and G3 regions (comprising the D’D3 and A1 domains, see Fig. 1), while stabilizing its overall topology. Strikingly, the S764-V2191 distance shows variant-dependent huge changes: 108.6 Å (p.R1205L), 194.9 Å (p.R1205C), versus 189 Å (WT). R1205 variant-induced conformational changes expose neo-epitopes, ultimately modifying VWF binding energetics to LRP1 domain IV. In fact, p.R1205L and p.R1205S bind to both the G2 and G1 regions, while WT, p.R1205H, and p.R1205C bind simultaneously to both G2 and G3 regions (see Fig. 4). Notably, both p.R1205C and p.R1205S were shown to be cleared even faster than WT-VWF in mice [3]. It is likely that phylogenetic pressure has selected an arginine residue at position 1205 to counteract excessive binding to macrophage receptors responsible for VWF clearance, thereby prolonging its persistence in the bloodstream.

As to the interaction with the platelet receptor GpIbα, the R1205X variants exhibit a significant decrease of their affinity for the receptor. Relative to WT-VWF, we observed 33-fold less affinity for p.R1205L/C variants, with more moderate reductions for p.R1205H (3-fold) and p.R1205S (10-fold). The p.R1205H variant is distinctive for retaining the highest residual GPIb affinity among VWF R1205X variants. Originally classified as “platelet-normal VWD1” [41], Vicenza-type VWD exhibits a unique phenotype: severe plasma VWF deficiency with normal VWF platelet stores, preserved large multimers [42] but defective RIPA, hallmarks that prompted its reclassification as type 2M. However, the clinical features of VWF “Vicenza” are still debated, because other studies found a normal or just slightly reduced functionality of the p.R1205H variant for platelets [42]. Regarding the interaction of p.R1205C/L with platelet GPIb, the EAHAD Coagulation Factor Variant Database [43] lacks specific data on potential impaired binding to GPIb and currently classifies these variants merely as “likely pathogenic”. Similarly, beyond its accelerated clearance from circulation, no data are available regarding the p.R1205S mutant’s interaction with GPIb [3]. From a structural viewpoint, the α1-β2 loop in the VWF A1 domain (residues L1309-V1314) plays a fundamental role for binding to platelet GpIbα [44]. In WT-VWF, complex formation with GpIbα causes the α1-β2 loop to rotate away from GpIbα and shift by over 6 Å [44]. In unliganded A1, the loop sterically clashes with GpIbα, blocking binding. The p.R1205X mutations, and especially p.R1205L and p.R1205C, disrupt this motion by destabilizing the α1-β2 loop (Figure 7, p.R1205L), thus preventing key interactions with K1332/R1334 and E225/D235 of GpIbα. Regarding the interaction of p.R1205C/L with platelet GPIb, the EAHAD Coagulation Factor Variant Database lacks specific data on potential impaired binding to GPIb and currently classifies these variants merely as “likely pathogenic”. The molecular modeling predictions and binding energy calculations presented in this study provide mechanistic insights that may help explain the clinical manifestations observed in VWD patients expressing these variants. This is the first in silico study that unravels the role of R1205 in modulating the mechanisms of VWF clearance and the interaction with the platelet receptor. These results could be relevant while designing and tailoring therapeutics in some types of VWD.

In summary, our modeling investigation shows that R1205 in VWF’s D3 domain serves two critical functions: a) modulating LRP1 domain IV binding and thus the protein’s clearance and b) maintaining a conformational state competent for GPIbα engagement. These findings underscore how R1205-driven structural dynamics dictate VWF function, a paradigm disrupted by pathogenic variants at this position. These findings provide a basis for experimental validation to verify the predicted biochemical interactions and determine also the precise subtypes of these natural VWF variants.

## Materials and Methods

### In-silico modeling of the missense VWF monomers and domain IV of LRP1

Arginine-1205 (R1205) is localized in the D3 domain of VWF, whose X-ray structure was solved and deposited in the PDB website along with other domains of the mature protein: D’D3 (S764-P1241, PDB file “6N29”), A1 (D1261-1468, PDB file “1AUQ”), A2 (M1495-H1674, PDB file “3GXB”) and A3 domains (D1685-S1873, PDB file “1ATZ”). Thus, we used the above X-ray-solved structures as templates, performing a molecular modeling of the stretched remaining sequences of the entire VWF region S1-V1428 (S764-V2191 for the full-length numbering system). Much progress has been made in protein structure prediction because of decades of effort [45–51]. Hence, the analysis of WT and pathological variants of VWF at R1205 was carried out using the I-TASSER threading modeling server (Iterative ThreadingASSEmbly Refinement) (http://zhanglab.ccmb.med.umich.edu/I-TASSER/), as previously detailed [52, 53]. The program generated multiple models, all exhibiting an elongated VWF conformation that was consistent across species. This elongation broadly matched both the extended cryo-EM structures of VWF tubules and the shear force-induced uneven protomer extension observed in atomic force microscopy studies. [54, 55]. Subsequently, each model was refined using the FG-MD program [56]. This program first identifies analogous fragments from the PDB by the structural alignment program TM-align [28]. Spatial restraints extracted from the fragments are then used to re-shape the funnel of the MD energy landscape and guide the molecular dynamics conformational sampling. Each molecular model of any VWF species was chosen only if TM-score was ≥0.5. This parameter is a proposed scale for measuring the structural similarity between two structures (in this case the predicted and the native structure of the solved VWF domains) [57]. Because the Root Mean Square Deviation (RMSD) is an average distance of all residue pairs in two structures, a local error (e.g., a misorientation of the tail) will raise a big RMSD value although the global topology is correct. In TM-score, however, the small distance is weighed stronger than the big distance which makes the score insensitive to the local modeling error. Hence, a TM-score ≥0.5 indicates a model of correct topology and a TM-score<0.17 means a random similarity. It has to be remarked that these cutoff values does not depends on the protein length [57]. FG-MD program aims at refining the initial models closer to the native structure. It can also improve the local geometry of the structures by removing the steric clashes and improving the torsion angle and the hydrogen-binding networks.

LRP1is a high molecular weight endocytic scavenger receptor that is highly expressed on cells of different tissues including macrophages [58]. The extracellular domain of LRP1 is structured on a modular structure consisting of four clusters (I, II, III and IV) of LDL receptor type A repeats that drive the interaction with several ligands [59]. In vitro studies have shown that cluster IV of LRP1 can bind to VWF [4, 60] and that LRP1 modulates VWF endocytosis into early endosomes [59]. The binding of wild type VWF to LRP1 occurs only when the former is in the stretched conformation, induced by high shear stress or ristocetin [39, 59]. No validated X-ray diffraction, NMR-based and cryo-EM structure of LRP1 is present in the PDB database. Hence, the extracellular CCR IV (Cluster IV of complement-like receptor) domain (cluster IV) of LRP1 (residues V3331-T3779) was modeled and refined by using the I-TASSER and FG-MD programs, as described above for the VWF monomer species.

### Molecular modeling of interaction between the VWF species and both LRP1 and GpIbα

Once both the VWF species and LRP1 Cluster IV models were refined, their docking complexes was investigated *in-silico* using the High Ambiguity Driven protein-protein DOCKing (HADDOCK) program version: 2.4 (https://rascar.science.uu.nl/HADDOCK2.4/) [61]. HADDOCK employs data-driven approaches [62], which integrate information derived from biochemical, biophysical or bioinformatics methods to enhance sampling, scoring or both [63]. The main characteristics of HADDOCK are the Ambiguous Interaction Restraints or AIRs, which allow the conversion of raw data including mutagenesis experiments or NMR chemical shift perturbation into distance restraints which are integrated in the energy functions that are used in calculations. In the docking protocol of HADDOCK, molecules pass through varying degrees of flexibility and distinct chemical surroundings. The performance of HADDOCK protocol depends on the number of models generated at each step. The grading of the clusters is based on the average score of the top four members of each cluster. The score is calculated as:

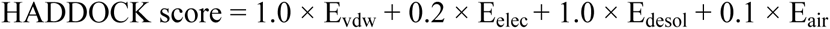

where, “E_vdw_” represents the intermolecular van der Waals energy, “E_elec_” is the intermolecular electrostatic energy, “E_desol_” is an empirical desolvation energy and E_air_ represents the AIR energy [64]. The cluster’s magnitude is indicated by its numbering in the results. On the results page, each cluster’s various components of the HADDOCK score are further explained. HADDOCK indicates that the top cluster is the most dependable. Better structures and interaction are indicated by higher negative HADDOCK and Z scores. Beginning with the individual structures of their constituent parts, HADDOCK is an integrative modeling technique that can integrate experimental and predicted data to direct the structure prediction of biomolecular complexes, such as protein–protein and protein–ligand complexes. HADDOCK drives the docking process via ambiguous interaction restrictions [61].

As to the interactions between the WT and the VWF variants with the N-Terminal domain of the platelet GpIbα, we used the best I-TASSER models of each VWF form and the X-ray diffraction form of GpIbα, in which structure of the N-Terminal region H1-R290 of the protein was solved at 1.7 Å (PDB DOI: https://doi.org/10.2210/pdb1P9A/pdb). Furthermore, the domains involved in the docking process and the relative contribution of hydrogen and hydrophobic energies were analyzed for each complex by using the PDBsum platform (http://www.ebi.ac.uk/thornton-srv/databases/cgi-bin/pdbsum/GetPage.pl?pdbcode=index.html). The graphical rendering of the various models was performed using the PyMol program.

### Energetics of WT and VWF variants models with both LRP1 Cluster IV and N-terminal domain of GpIbα

The prediction of the binding affinity between Cluster IV_LRP1-VWF models and GpIbα-VWF models was calculated by using the PRODIGY program, which quantifies the interaction energy of protein-protein complexes from their 3D structure, taking into consideration hydrogen and hydrophobic contributions to binding energy [65].

## Author contributions

MS and SL: data curation (equal); methodology (equal); visualization (equal); writing – review and editing; AF, MB, and LDG, critically reviewed the manuscript, GC and RDC: conceptualization (equal); funding acquisition (RDC); methodology (RDC); supervision (equal); writing – original draft (RDC); writing – review and editing (equal).

## Acknowledgements

RDC gratefully acknowledges funding from the Catholic University of the Sacred Heart, School of Medicine (Linea D1).

